# Inter-ecosystem variation in the food-collection behaviour in climbing perch *Anabas testudineus*, a freshwater fish

**DOI:** 10.1101/573600

**Authors:** V. V. Binoy, V. B. Rakesh, Anindya Sinha

**Author notes:** Corresponding author*: V.V. Binoy, National Institute of Advanced Studies, Indian Institute of Science Campus, Bangalore 560012, India.

## Abstract

A unique piscine behaviour—collection and temporary storage of food materials inside the mouth during times of availability and particularly in response to starvation—has been reported in a single species, the climbing perch *Anabas testudineus*. In this study, we documented a significant variation in the amount of food collected by populations of climbing perch inhabiting different ecological regimes, *kole* paddy fields, canals and water channels in coconut plantations, after experiencing starvation for two different periods of 24 and 48 h. Our results revealed a significant flexibility in this unique behaviour, depending on the ecological conditions and hunger experienced by the individuals.

## Introduction

The collection and storage of food for later consumption is an effective strategy displayed by different species for the successful exploitation of transitory food resources and to out-compete potential competitors while foraging in groups [1, 2]. Additionally, the ability to move food materials out of predation-prone feeding grounds to safer areas for consumption enhances the personal safety of the individual [3]. This effective foraging strategy of collection and stocking of food for either short or long durations of time was traditionally considered the domain of only some species of birds and mammals [2, 4] until Binoy and Thomas [5] added a freshwater fish, the climbing perch *Anabas testudineus*, to the list. This fish species, the only one known so far to display this unique behaviour, collects food materials in its mouth before ingestion, in a fashion possibly analogous to the storage of food in the cheek-pouches of cercopithecine primates [4]. Interestingly, as reported in various mammalian species, the climbing perch also enhances food collection over food consumption after experiencing food deprivation [5]. We would, however, like to point out that although this behaviour, as shown by the climbing perch, was described as ‘food stocking’ by Binoy and Thomas [5], such a term better applies to the relatively more prolonged periods of food storage shown by birds and mammals [2]. The term ‘food collection’ may be more appropriately used to denote this behaviour, as the climbing perch collects and stores food materials in its mouth for a few minutes alone [5].

The climbing perch (*Anabas testudineus* Bloch) is a shoal-living fish, inhabiting different kinds of lentic and lotic freshwater ecosystems in India and several southeast Asian countries [6]. Being equipped with the labyrinthine organ to breathe atmospheric air, this fish has been reported from waterbodies such as ponds, rivers, marshes, sewage canals, irrigation canals, *kole* paddy fields and areas of saline intrusion, all of which differ significantly in their ecological characteristics [7]. In natural habitats, this fish consumes a wide range of food items including crustaceans, worms, molluscs, insects, algae, soft parts of aquatic plants and organic debris.

It is a well-known fact that the tracing of modulations that characterise the vital components of the foraging behaviour of a species in relation to the changes happening in the biotic and abiotic components of its environment is crucial for a comprehensive understanding of the evolution of its foraging behaviour [8]. The climbing perch thus offers an excellent model system to study this vital topic in a piscine species due to its tremendous capacity to adapt to starkly contrasting ecological conditions [7].

Furthermore, such information will also be useful in optimising food consumption behaviour and hence, enhancing the survival and growth of this economically important fish in artificial environments in which it is cultivated extensively. Unfortunately, till date, very few studies have explored foraging behaviour of this species in natural or artificial environments [7, 9, 10], no information is available on the food collection behaviour of climbing perch populations surviving in contrasting ecological conditions or on the factors influencing this behaviour.

The current paper examines variation in food-collection behaviour of climbing perch individuals from three different ecosystems—*kole* paddy fields, shallow water channels in coconut plantations and canals—and deprived of food for two different durations, 24 and 48 h.

## Materials and Methods

### Subject fish and husbandry

Climbing perch were collected from the following three ecosystems, located in different parts of Thrissur district of Kerala state, southern India.

#### Water channels

These shallow channels are characteristic of coconut and banana plantations of central Kerala [7]. The climbing perch were collected from channels of a coconut plantation, located at Edathirinji (10.33°N, 76.18°E). This particular water body (mean water depth ± SD of 60 ± 20 cm and breadth, 50 ± 15 cm), situated in between the rows of coconut plants, was turbid, foul-smelling, due to decaying plant materials, and occupied by various aquatic and semi-aquatic plants. Although poor-quality water and reduced availability of space in this shallow ecosystem supported very few piscine species, the climbing perch survived in this extreme environment due to its capacity to breathe atmospheric air and pliantly available food materials, insect larvae and decaying plant materials [7].

#### Canals

This lotic ecosystem, from which the fish were collected, was located at Irinjalakuda (10.34°N, 76.19°E) and was contaminated by sewage from houses present on its banks. The climbing perch population living in this ecosystem had to face considerable harshness of water flow, transitory food materials as well as the reduced availability and diversity of food items [7].

#### Kole paddy fields

*Kole* is a unique agro-wetland ecosystem, located in the central region of Kerala state. Being positioned in the basins of two major rivers, this paddy field resembles a large shallow lake during the monsoons and is utilised for cultivating paddy for the rest of the year [11]. The *kole* paddy fields are typically extensive in distribution, hold good-quality water, rich in spatial complexity and exhibit higher levels of primary and secondary productivity [12], as compared to the irrigation canals and channels.

The fish were collected from the Irinjalakuda (10.35°N, 76.21°E) region of the *kole* and other two focal ecosystems with the help of expert fishermen and transferred to the laboratory. Only fish of standard length (mean ± SD of 62.4 ± 26.0 mm) were used in the experiments. Due to the lack of sexual dimorphism in this species [13], however, the subject fish could not be sexed.

Individuals from each ecosystem were kept isolated in aquaria (45 × 22 × 22 cm) and these aquaria themselves were used for the subsequent experiments involving the subject fish. Water temperature was maintained at 25 ± 1°C and light hours at 12L: 12D. Three sides of all aquaria were covered with black paper while steel grids, placed on the top, prevented the fish from jumping out. Twenty-five food pellets were dropped together at a specific site in the aquarium once a day (between 09:00 and 10:30) to acclimatise the fish with a specific feeding schedule [5]. No hesitation was shown by the subject fish in consuming the commercial food pellets from the first day of isolation itself. Unused pellets were siphoned out 30 min after their addition to the aquarium.

After giving five days for acclimating with the laboratory environment, the food-collection behaviour was tested as follows. In order to standardise their hunger state, the subject fish were deprived of food materials for 24 h before the experiment. Food pellets of length (mean ± SE) of 3.58 ± 0.14 mm and mass 0.01 ± 0.003 g were dropped, one by one, at intervals of 1 s at the same site in the aquarium. The subject fish continuously collected the pellets in their mouth (see also Binoy and Thomas 2008 and Supplementary Material S1). Pellets falling out of the mouth of the fish due to overfilling or the individual not gathering food granules for a continuous period of 120 s were marked the end points of the experiment. The same fish were re-tested to evaluate the impact of prolongation of the starvation period to 48 h, six days after the first experiment, by following the same protocol^5^. The availability of climbing perch was much less in the water channel and canal ecosystems, in comparison to the *kole* paddy fields and hence, 16 individuals were tested from each of the first two ecosystems and 20 fish from the *kole* paddy field.

The parametric tests ANOVA, Tukey’s test and paired *t*-test were used for analysis as the data was established to follow normal distribution (Kolmogorov-Smirnov Test).

## Results

The study climbing perch populations, collected from different ecological conditions, differed significantly in their ability to collect food materials inside their mouths after experiencing food deprivation for 24 h (ANOVA, *F_2,49_*= 4.63, *p* < 0.05; Fig. 1). The fishes from the canal ecosystem collected significantly greater number of pellets than did their counterparts from the water channels (Tukey’s test, *tij* = −3.13, *p* < 0.01). However, no significant difference was observed in the number of pellets collected by the members of the canal and *kole* paddy field (*tij* = −1.93, *p* > 0.05) or between the water channel and *kole* paddy field populations (*tij* = 0.95, *p* > 0.05).

**Figure 1.**
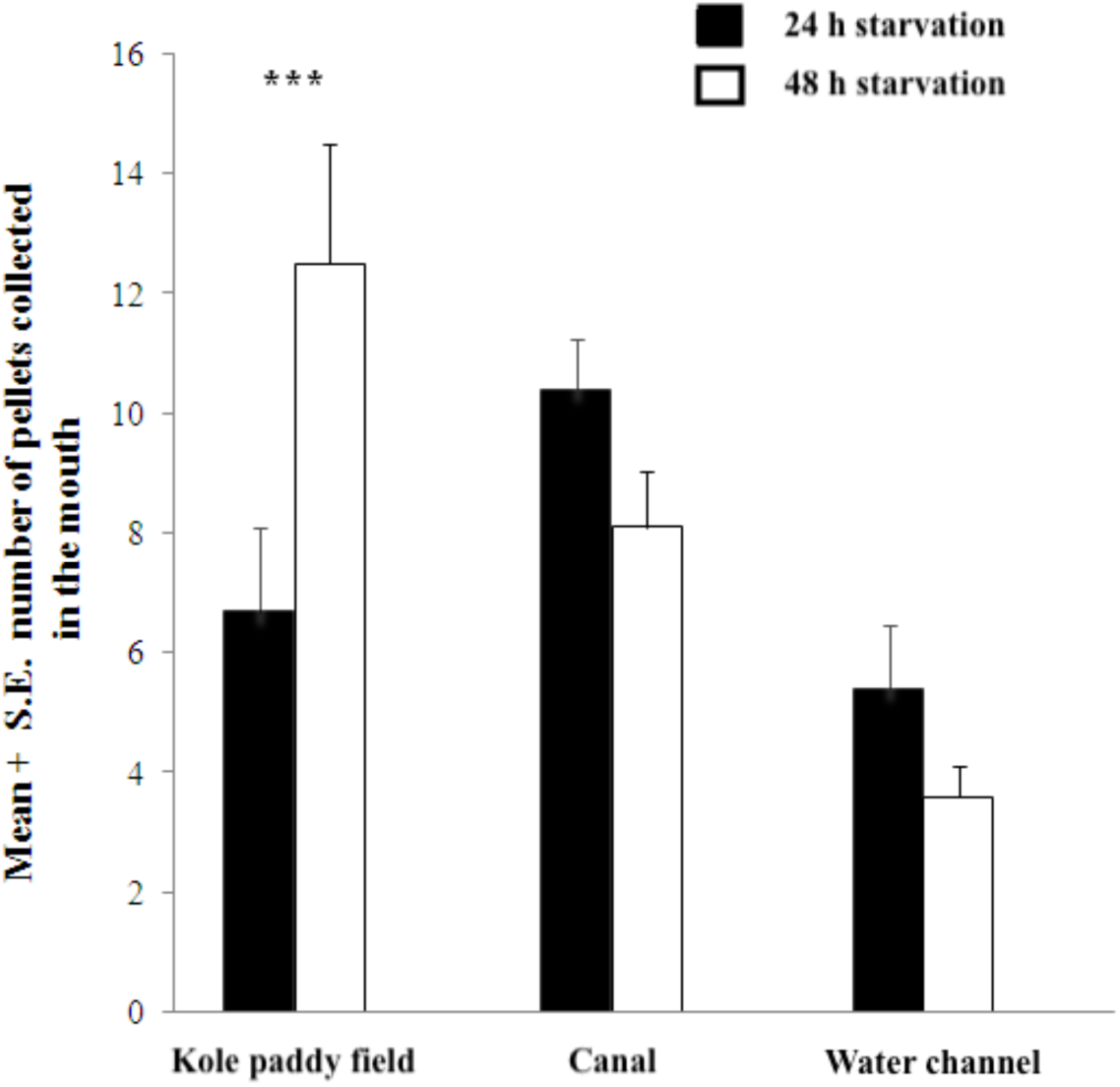
Mean number of food pellets collected in the mouth by climbing perch living in different ecosystems, after experiencing food deprivation for 24 and 48 h.

Only individual climbing perches from the *kole* paddy field exhibited significant modification in the food collection behaviour (ANOVA, *F_2,57_* = 9.26, *p* < 0.001) when the duration of food deprivation was increased from 24 to 48 h. These fish collected significantly more food material in the mouth in response to prolonged starvation than did their counterparts from the canals (*tij* = 2.55, *p* < 0.05) or from water channels (*tij* = 4.79, *p* < 0.001). However, channel and canal populations were not significantly different from one another in the performance of this unique behaviour when the duration of food derivation was doubled (*tij* = −2.12, *p* > 0.05).

Differences in the amount of food pellets collected by the individuals from each of the three focal populations, after experiencing starvation for the two different periods, was also analysed. The climbing perch from *kole* paddy fields almost doubled the amount of food collected (paired *t*-test, *t_19_* = −4.17, *p* < 0.001; Fig. 1) when the food deprivation was increased from 24 to 48 h but no such behavioural modification was observed in the individuals from irrigation canals (*t_15_* = 2.12, *p* > 0.05) or from water channels (*t_15_* = 1.74, *p* > 0.05).

## Discussion

The marked differences in the food-collection behaviour of the climbing perch, collected from three different ecological conditions, could be the result of variation in the food availability, inter-and intra-specific competition and predation pressures experienced by the individual fishes in their respective habitats [1,5]. The climbing perch living in the canal, possibly the harshest amongst the three ecosystems studied, have to survive on the very little food material available in their surroundings. The prevalent turbidity and water flow could also increase the difficulty of food acquisition manifold for this visually orienting species [14] in the canals. According to Binoy and Prasanth [7], the only food present in the gut of the climbing perch, collected from this ecosystem, were insect larvae (mainly *Chironomus* spp.) and debris, and the studied individuals had rather low amounts of food in their digestive tracts. Hence, the climbing perch from such an ecosystem could be collecting the maximum number of food pellets it could, at the time of availability, which reflected as significant deviation from those taken by the fish from the water channel and *kole* paddy field populations.

In contrast to the canal and water channel ecosystems, the *kole* paddy field constitutes a geographically widespread habitat covering more than 10 thousand hectares and famous for the diversity of microhabitats and life forms that it harbours [12]. However, the climbing perch inhabiting this ecosystem typically have to face increased levels of both predation pressures and competition for food materials from conspecific as well as heterospecific individuals, in comparison to their counterparts from other two focal habitats. Generally, air-gulping piscine species synchronise their surfacing activity in order to reduce the probability of falling prey to aerial predators [15]. Accordingly, the number of climbing perch observed preforming orchestrated air-gulping activity in each bout was considerably high in the *kole* paddy fields (pers. obs.).

Moreover, an exploration of the catch of fishermen revealed that the number of individuals of the major piscine predators of the subject species, the snakeheads (*Channa striatus* and *C. marulius*), were much less prevalent in the water channel and canal ecosystems (pers. obs.). Enhancing levels of food collection, therefore, could be a behavioural adaptation exhibited by hungry climbing perches living in the *kole* paddy field ecosystem, to out-compete their conspecifics or heterospecifics by acquiring more food during times of availability. The climbing perch investigated in a previous study [5] were captured from a large pond, which resembled a *kole* paddy field in its ecological properties, and showed a similar enhancement in their food collection strategy. Furthermore, a recent study by Zworykin [10] has proven that the presence of conspecific individuals could influence the feeding behaviour of climbing perch; this study reported a significant variation in the foraging behaviour of individuals when tested in isolation and in presence of the conspecifics. It may also be noted here that a similar analogous enhancement of food stocking in the cheek pouch to out-compete competitive conspecifics during intensive feeding bouts has also been reported in troops of different cercopithecine primate species [1, 5].

In conclusion, therefore, individuals of species such as the climbing perch could vary their ability to utilise food resources due to their adaptation to the food web in which they are embedded [16, 17]. Additionally, however, the species that we studied is famous not only for its specific ability to survive under a wide range of ecological conditions but also for migration over land and consequently changing habitats during the monsoons [18]. An earlier study by our group [7] revealed that climbing perch populations living in contrasting ecological conditions display great disparity in the nature and quantity of food items consumed. A detailed analysis of the adaptive modifications of foraging behaviour by this species in dissimilar ecosystems could thus reveal the interplay between physiological, social and environmental factors in determining the unique food-collection behaviour displayed by this species.

## Acknowledgements

VVB is grateful to the Science and Engineering Research Board (SERB), Department of Science and Technology, Government of India, for a Young Scientist (Fast Track) Grant (SB/FT/LS-155/2012) that enabled this study. The experiments reported in this paper comply with the current relevant laws of India.

## Legend to the Electronic Supplementary Material S1

The study species, climbing perch, collecting food pellets for storage in its mouth.

